# Mechanical morphogenesis and the development of neocortical organisation

**DOI:** 10.1101/021311

**Authors:** Ophélie Foubet, Roberto Toro

## Abstract

The development and evolution of complex neocortical organisations is thought to result from the interaction of genetic and activity-dependent processes. Here we propose that a third type of process – mechanical morphogenesis – may also play an important role. We review recent theoretical and experimental results in non-linear physics showing how homogeneous growth can produce a rich variety of forms, in particular neocortical folding. The mechanical instabilities that produce these forms also induce heterogeneous patterns of stress at the scale of the organ. We review the evidence showing how these stresses can influence cell proliferation, migration and apoptosis, cell differentiation and shape, migration and axonal guidance, and could thus be able to influence regional neocortical identity and connectivity.

## 1. Introduction

The neocortex is the most distinctive structure of the mammalian nervous system. Whereas its thickness and radial organisation vary little among species, its surface area can display >1000-fold changes, increasing the number of its specialised regions and the complexity of its connectivity. This increase in complexity is the most likely biological substrate of the increased behavioural sophistication that leads to the emergence of cognitive functions such as consciousness, creativity and language. How do complex neocortical organisations develop and evolve?

Today, neocortical organisation is thought to result from the interaction of genetic and activity-dependent processes. During development, morphogens and signalling molecules secreted from a reduced number of patterning centres are thought to produce gradients of expression of transcription factors, which together with neuronal activity driven by thalamo-cortical afferents establish and maintain areal identity (O’Leary et al., 2013). From the perspective of evolution, an increase in neocortical complexity should result from the increased sophistication of the genetic program that would control areal identity directly, or indirectly, by first establishing the connections that will later enable activity-dependent processes to operate.

Here we propose that a third type of process – mechanical morphogenesis – may also play an important role. Theoretical and experimental results from the physics of soft tissues show that homogeneous growth can produce a rich variety of forms. In particular, several biophysical models suggest that neocortical folding could be caused by this type of growth-induced mechanical instabilities. The resulting forms are associated with complex patterns of mechanical stress at different scales. The developing tissue is very sensitive to these mechanical signals, which can influence cell proliferation, apoptosis, cell differentiation, cell shape, cell migration and axonal guidance. Through mechanical morphogenesis, a broad regulation of growth could then lead to an increased number of heterogeneous areas, and the macroscopic changes in neocortical geometry could facilitate the formation of new connectivity patterns.

We will first summarise the role of activity-driven and genetic processes in the production of neocortical arealisation and connectivity. We will then address the problem of explaining the relationship between neocortical arealisation and folding. To better understand neocortical folding and mechanical morphogenesis, we will briefly introduce the theory of mechanical elasticity and growth. A website with interactive mechanical simulations illustrating these principles can be accessed at http://neuroanatomy.github.io/growth. Finally, we review recent results highlighting the effect of mechanics on the developing brain tissue and discuss a mechanical morphogenesis hypothesis for the development and evolution of neocortical organisation, its predictions, and experimental approaches that should allow us to test them.

## 2. Development of neocortical arealisation

> *“It must remain an open question whether the refinement of the cortex through differentiation is always the result of external, physical causes, or whether many of the associated phenomena may be explained in other ways, unrelated to external living conditions and unrelated to the struggle for existence, rather due to a property of the organism itself, an ‘energy for refinement’ (R. Hertwig) or, as Naegli expresses it, a ‘principle of progression’“* (Brodmann, 1909; Brodmann and Gary, 2006).

The neocortex is the outermost part of the cerebral hemispheres. In the radial direction it is composed of 6 main layers containing neuronal cell bodies: the grey matter. The myelinated axons of these neurones constitute most of the cerebral white matter. Together, the neocortical grey matter and the white matter represent ∼90% of the human brain. The thickness of the neocortex varies little among species. The neocortex of mice has an average thickness of 1.5 mm, compared with 2.5 mm in humans. By contrast, there are extreme differences in cortical surface area among mammals, a 1000-fold difference between mice and men. The expansion of the neocortex correlates with the development of cortical folds and the formation of an increased number of specialised regions, the cortical areas. Based on cytoarchitecture, chemoarchitecture and connectivity patterns, it is possible to distinguish ∼20 cortical areas in lissencephalic species such as mice, but ∼50 in gyrencephalic species such as macaques, and probably more than 200 in humans (Kaas, 2012). In gyrencephalic species, multiple studies have revealed a close relationship between cortical folding and the cytoarchitectonic, connective and functional organisation of the neocortex, suggesting that common biological processes shape them. For example, in racoons the folds of the somatosensoty cortex precisely separate the functional fields of the palm and the fingers (Welker and Campos, 1963; Welker et al., 1990). In primates, the central sulcus divides the primary motor cortex from the somatosensory cortex, and the calcarine sulcus divides the superior and inferior visual hemifields of the primary visual cortex in the occipital lobe (Brodmann, 1909; Brodmann and Gary, 2006; Fischl, 2013; Fischl et al., 2008).

The major divisions of the nervous system develop during the early segmentation of the neural tube into neuromeres (Puelles, 2009, 2013). The neocortex is essentially a homogeneous structure – an isocortex – which develops from the rostral-most neuromere. Only a few cortical areas, such as the primary visual cortex, can be distinguished with the naked eye. For the most part, it was only after the development of staining and tracing techniques that neuroanatomist were able to describe them. A core group of areas is present in all mammals. These areas, the primary sensory and motor cortices, receive preferential innervation from specific thalamic nuclei. In species with large cortices there is a progressive addition of multi-modal associative areas, connected mostly with other neocortical regions but also with thalamic nuclei (Krubitzer, 2009).

Despite their stability within a single species, neocortical areas are dynamic entities, whose organisation can be dramatically modified by changes in neuronal activity. For example, in animals with stereoscopic vision such as cats or monkeys, the primary visual cortex receives thalamic afferents associated with the left and the right eyes clustered into bands of ocular dominance. Early visual deprivation can change the balance between left and right innervation, or even completely suppress the formation of bands (Espinosa and Stryker, 2012). The processes necessary to dynamically develop this type of cortical modules seem to be shared by all vertebrates, and ocular dominance bands can be made to develop even in species which do not have them naturally. For example, the primary visual cortex of mice, or the optic tectum of frogs, are innervated by axons related to the contra-lateral eye only, and do not develop ocular dominance bands. Bands can be made to develop, however, by driving connections from another eye, either by altering axonal guidance through gene knockout in mice (Merlin et al., 2013) or by grafting a 3rd eye in frogs (Constantine-Paton and Law, 1978). The link between primary cortices and specific sensory modalities can be also modified by activity. In a series of experiments in ferrets, Sur and colleagues (Roe et al., 1990, 1992; Sharma et al., 2000; Sur et al., 1988) forced visual afferents to innervate what would normally be the auditory cortex, making it acquire many of the cytoarchitectonic and functional properties of the visual cortex (Sharma et al., 2000). Despite using their auditory cortex for vision, rewired ferrets exhibited visual behaviour similar to that of normal ferrets (von Melchner et al., 2000).

Because of this remarkable plasticity, neuronal activity was originally thought to be the primary cause of neocortical arealisation. The existence of intrinsic processes, however, had been hypothesised since the first treatises on cortical cytoarchitecture (Brodmann, 1909). The idea was much discussed at the end of the 80s with the introduction of the protomap (Rakic, 1988) and the protocortex (O’Leary, 1989) hypotheses, which emphasised differently the role of genetic and activity-dependent processes in neocortical arealisation. The first direct evidence for a genetic process came 10 years later, with the demonstration of the roles of Emx2 and Pax6 in the specification of cortical progenitor identity in mice (Bishop et al (2000), Mallamaci et al (2000), see O’Leary et al. (2013) for a review). Subsequent studies showed the existence of several patterning centres located at the periphery of the neocortex, secreting morphogenes and signalling molecules such as members of the fibroblast growth factor family (FGFs) and bone morphogenetic proteins (BMPs). These molecules formed gradients over the ventricular zone – where most neocortical neurones are produced in mice – which controlled the expression of transcription factors such as Emx2, Pax6, COUP-TF1, or Sp8 in cortical progenitors. Neurones produced in the ventricular zone migrate radially to form the cortical plate, which later matures into the neocortex. Each position of the ventricular zone could be then characterised by a unique combination of transcription factors, encoding the neocortical areal identity that is propagated through radial migration. The experimental alteration of the morphogenetic gradients, or the expression of transcription factors has been shown to result in the displacement or even the duplication of cortical areas (Fukuchi-Shimogori and Grove, 2001), as well as marked changes in the proportions of others (Hamasaki et al., 2004).

Today, intrinsic genetic and extrinsic activity-dependent processes are thought to act in combination to produce neocortical arealisation. The initial gradients of gene expression in the ventricular zone and the cortical plate would be genetically encoded, defining in some cases areal identity directly, or defining first the connectivity patterns that would later allow neuronal activity to refine the basic area map, producing the discrete areas and modules of the adult neocortex (O’Leary et al., 2013). The phylogenetic differences in neocortical complexity between mice and humans, for example, would result from differences in the sophistication of the genetic program that either prescribes areal identity directly, or indirectly by encoding first connectivity.

But how to explain the relationship between neocortical arealisation and brain folding in gyrencephalic species? One possibility is that the same genetic program that encodes arealisation (directly, or indirectly through neuronal activity) would guide the formation of folds at specific locations (Ronan and Fletcher, 2014). Species-specific folding patterns would then be a reflection of the genetic programs that produce the different neocortical area maps. For example, gyri could result from bulging due to a locally higher production of neurones (Borrell and Götz, 2014; de Juan Romero et al., 2015; Nonaka-Kinoshita et al., 2013; Reillo et al., 2011; Ronan and Fletcher, 2014; Smart and McSherry, 1986a, 1986b; Stahl et al., 2013; Welker et al., 1990). Borrell and colleagues have reported that in ferrets the density of dividing progenitors at P3 (3rd postnatal day, when the ferret cortex is still smooth) was higher under the region that will become later the splenial gyrus compared with the neighbouring regions, that will become sulci. Alternatively, folds could result from bending due to the active contraction of axonal fibres connecting different neocortical areas together (Van Essen, 1997; Geng et al., 2009; Hilgetag and Barbas, 2005, 2006). Van Essen (1997) reported that neighbouring areas with strong interconnections were more often on facing sides of a gyrus, compared with weakly connected areas, which tended to be separated by a sulcus. In all cases, the same genetic program that encodes brain connectivity would produce neocortical folding as a by-product.

However, these explanations do not take into account the physics of growth and the geometric and mechanical properties of the developing brain. The mechanical stress patterns that would produce brain folding if it were due to bulging or bending are not compatible with those observed in real brains (Xu et al., 2009, 2010). Recent theoretical and experimental evidence suggest that brain folding is more likely the result of a buckling instability induced by global neocortical growth: submitted to homogeneous growth, physical systems similar to the developing brain produce folding patterns strikingly reminiscent to those of real brains. The intrinsic “energy for refinement” alluded by Brodmann could well be both biological and mechanical.

## 3. The physics of mechanical morphogenesis

### 3.1. Elasticity

Brain tissue is composed in a large part of water, it is mostly incompressible (its volume is conserved despite deformation), elastic (after deformation it recovers its original shape), and to some extent plastic (strong or prolonged deformation produce a permanent change in shape). The elastic forces produced by deformation can be calculated relative to an idealised rest configuration, where the tissue is free of mechanical stress (see Fig. 1). Imagine that we subdivide the tissue at rest into small volume elements, and then that we look at how each of them is transformed in the deformed configuration (Fig. 1a). Each volume element can be described by a small cube, and a corresponding stretched and skewed box in the deformed tissue (Fig. 1b). It is possible now, for each volume element, to compute a 3×3 transformation matrix **F** that changes the cube at rest into the deformed box (Fig. 1c-e). This transformation matrix **F** is called the *deformation tensor*.

**Figure 1.**
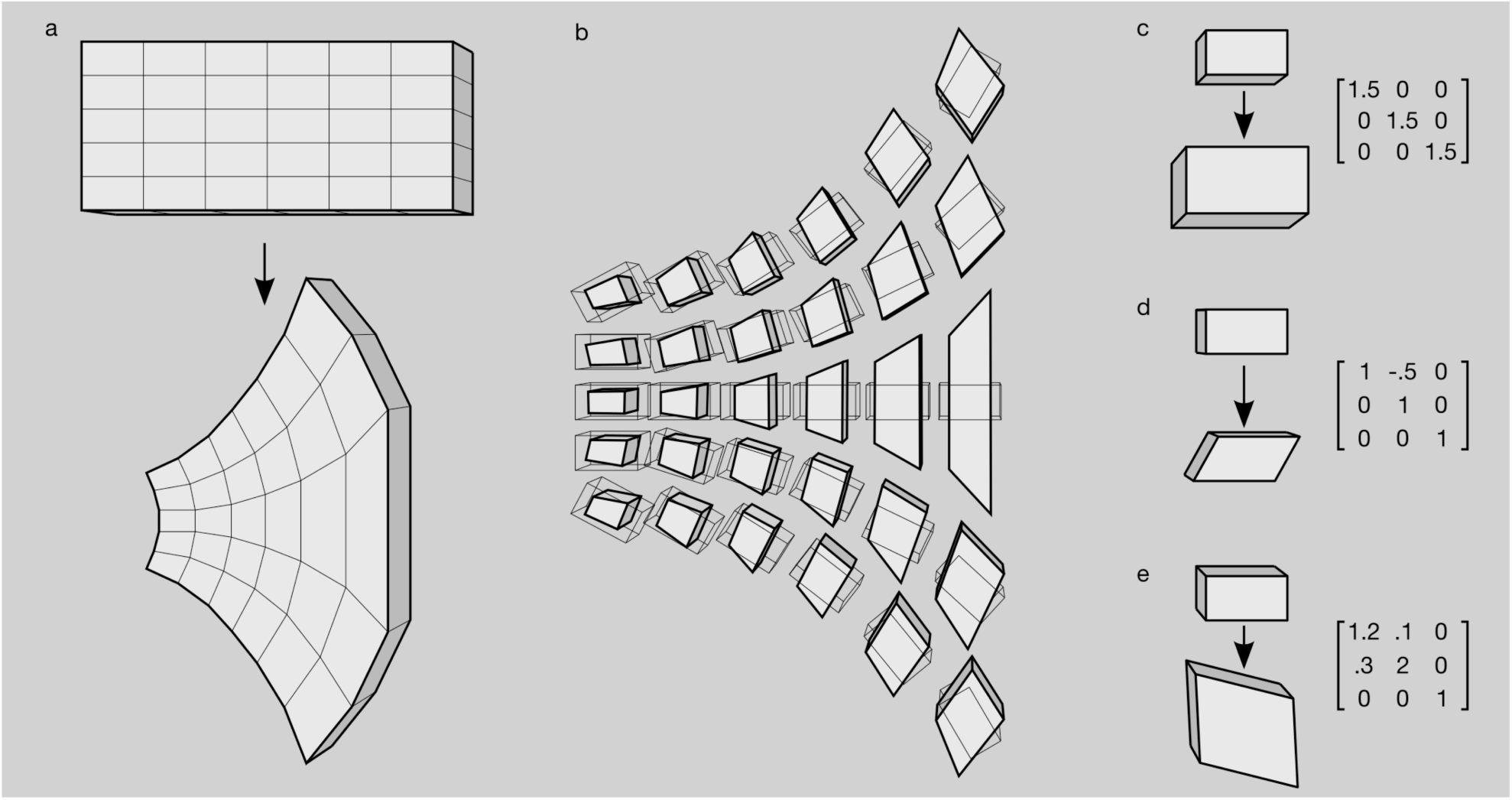
**Deformation and elasticity**. Tissues arc decomposed into a continuous grid of volume elements. The elastic deformation of each volume element is measured relative to an idealised resting configuration. The deformation of each volume element is described by a matrix, the deformation tensor, (a) Deformation of a tissue. The tissue is decomposed into a continuous grid of volume elements, (b) The deformation of each volume element is measured with respect to the shape that it would ideally have at rest, (c) Matrix representation of a change in volume without change in shape, (d) Matrix representation of change in shape without change in volume, (e) Matrix representation of a combined change in shape and volume.

In order to compute the mechanical forces produced by deformation, we need a *constitutive model* that will describe how deformation and force are related. One of the simplest constitutive models is Hook’s law, which states that force is proportional to deformation. In the case of a linear spring, for example, elastic force would be directly proportional to elongation. The proportionality constant is known as *Young’s modulus* (*E*) whose unit of measurement is the Pascal (Pa). Brain tissue is very elastic, similar to jelly, *E* ∼1.5 kPa. By contrast, Young’s modulus for rubber is *E* ∼10-100 kPa, and for metal *E* >10 GPa.

In two or three dimensions, a new parameter is required to describe how much pulling in one direction will produce stretch in the orthogonal directions: *Poisson’s ratio* (ν). Stretching a square of rubber to twice its length, will make it shrink to half its width: a Poisson’s ratio of 0.5. By contrast, compressing a piece of cork in one direction produces almost no change in the others: the Poisson’s ratio of cork is close to 0. A linear constitutive model will include these 2 constants *E* and ν It is sometimes convenient to combine Young’s modulus and Poisson’s ratio into a bulk modulus *K* and a shear modulus μ. The bulk modulus represents the amount of mechanical stress produced by changes in volume, whereas the shear modulus represents the amount of stress produced by changes that do not affect the element’s volume.

But biological tissues can be described using a linear constitutive model only for very small deformations. Most frequently, a hyperelastic constitutive model is required, where the relationship between deformation and force is non-linear. A widely used hyperelastic constitutive model is the Neo-Hookean model introduced by Rivlin (1948). The model is defined in terms of the elastic energy *W* that the deformation described by the tensor **F** will produce in a material of shear modulus μ and bulk modulus *K*:

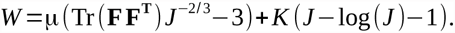

In this equation *J* measures the change in volume produced by the deformation **F** (mathematically, *J* =*det* (**F**)). If a volume element is shrunk by **F** then 0< *J* <1, if it is expanded, then *J* >1, and if volume is conserved as in the case of incompressible materials, then *J* =1. The right-hand side of the equation has 2 components. The first one, multiplied by μ, increases the energy *W* as the shear components of **F** increase (the off-diagonal elements of the matrix **F**). The second part, multiplied by *K*, increases *W* as **F** produces volume changes. In this second part, if there is no change in volume then *J* =1, log (*J*)=0, and there is no change in energy due to bulk volume changes. If **F** shrinks the volume element so that *J* is close to 0, then-log (*J*) will be very large, increasing the total elastic energy *W*. *W* will also increase if *J* >1, but the increase will not be proportional to the deformation because of the subtraction of log (*J*) : in the Neo-Hookean model disproportionately larger forces are required to keep increasing the deformation.

### 3.2. Growth

Growth introduces a different kind of deformation. In an influential article, Rodriguez et al (1994) conceptualised the deformation induced by growth as the combination of a term related to elasticity and another related to growth itself:

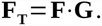

Applying the deformation tensor **F_T_** to a volume element is then equivalent to make it grow first, and introduce the deformation due to elasticity next. Only the elastic part of the deformation, however, changes the elastic energy *W*.

Consider for example a ring of elastic material that grows only in circumference but not in thickness. If we cut the ring in two pieces before it grows, as shown in Fig. 2, the pieces could be stuck back without introducing any force. After growing, however, the length of the ring will have increased so that sticking the pieces back will introduce a residual stress all over it. After sticking the pieces, the ring will undergo compression in the circumferential direction, and depending on its Poisson’s ratio, it will also undergo a force in the radial direction. Growth can be thought as a deformation of the resting configuration of the volume elements of an object, independently one from another. The requirement of continuity of the object will then introduce elastic forces, and a field of residual stress across the object (Fig. 2c).

**Figure 2.**
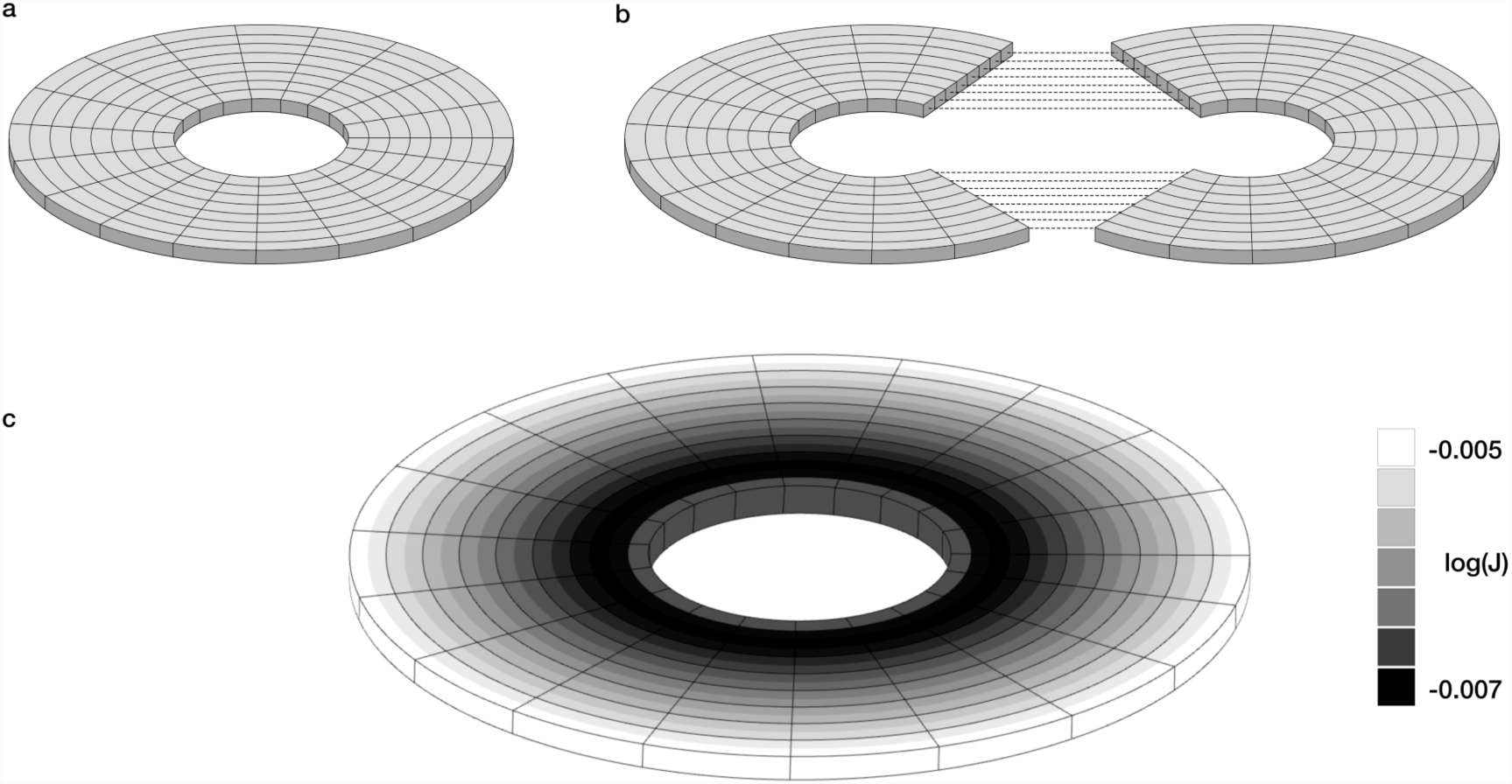
**Residual stress induced by growth.** Growth can lead to a configuration where stress cannot be eliminated, producing a gradient of residual stress. In this example, a ring originally at rest is made to grow such that its perimeter increases by 50% but without increasing neither the radial length of the ring nor its thickness. At equilibrium, the ring presents with a gradient of residual stress, (a) Original configuration of the ring at rest, (b) After growth, the ring could reach a zero stress configuration only if it were cut into two pieces. The dashed lines show the regions that should have to be stuck together to reconstruct the original ring, (c) The uncut ring after growth exhibits a gradient of residual stress. The ring is contracted overall, but especially within a band near its inner side. Grey level: logarithm of the deformation (Jacobian of the deformation tensor). Black: log(J)=-0.007, white: log(J)=-0.053, bulk modulus *K* =50, shear modulus *μ* =1. An interactive simulation illustrating the formation of a residual stress gradient due to growth can be accessed at http://neuroanatomv.github.io/growth

The morphogenetic effect of growth within the framework proposed by Rodriguez et al. (1994) has been addressed in several theoretical studies and confirmed experimentally. The elastic instabilities induced by growth can explain aspects of the shape of flowers (Amar et al., 2012), the formation of villi and crypts in intestinal epithelium (Hannezo et al., 2011), growing tumours (Ciarletta, 2013) or mucosa wall patterning (Xie et al., 2014) among many others. In particular, the effect of growth and elasticity in systems resembling the cerebral cortex has been observed to introduce different buckling instabilities which are able to produce folds of a geometry and a scale similar to those of real brains.

### 3.3. Buckling instabilities and neocortical folding

During neuronal migration from the ventricular zone (Rakic, 1988; Rakic et al., 2009) and the subventricular zone (Hansen et al., 2010; Reillo et al., 2011) the cerebral cortex of many gyrencephalic mammals is still smooth and mostly undifferentiated (Welker et al., 1990). Once cell migration is finished, the neocortex starts a period of fast expansion, due to the increase of intracortical connectivity, dendrites and glia.

At this early stage, the mechanical system composed by the neocortex and subcortical structures (including radial glia and the developing white matter) can be viewed as a system of two elastic substrates, one of which undergoes much more growth than the other. The deformation of an elastic layer whose surface area is much larger than its thickness (a *thin plate* in mechanics) is often described using Föppl-von Kármán equations. Although there is no general solution for these equations, they can be solved for particular cases or approximated through numerical simulation.

When submitted to load, shrinkage or growth, Föppl-von Kármán equations predict different types of buckling instabilities. For small growth, stretching is energetically less costly than bending and the system grows without buckling (this is likely the reason why small brains are lissencephalic). As growth increases, however, there is a critical threshold after which bending becomes less costly than further stretching and the system buckles. Among these buckling instabilities, wrinkles (Bowden et al., 1998; Cerda and Mahadevan, 2003; Davidovitch et al., 2011), folds (Pocivavsek et al., 2008) and creases (Hohlfeld and Mahadevan, 2011; Tallinen et al., 2014) can produce patterns such as those observed in gyrencephalic brains. Wrinkles develop when a stiff, thin, elastic layer – a skin – grows over a softer one. Wrinkles are sinusoidal undulations and are the first instabilities observed after buckling (see Fig. 3 for an illustration). They can develop into folds, where deformations are more localised, with flat regions coexisting with valleys or depressions of the stiff layer. Creases (also called sulci) are similar to folds, but they do not require a difference in stiffness between skin and substrate to develop. Whereas the depth of wrinkles and folds is smooth, similar to the first stages of gyrogenesis, the depth of creases is cusped, similar to the shape of adult brain sulci.

**Figure 3.**
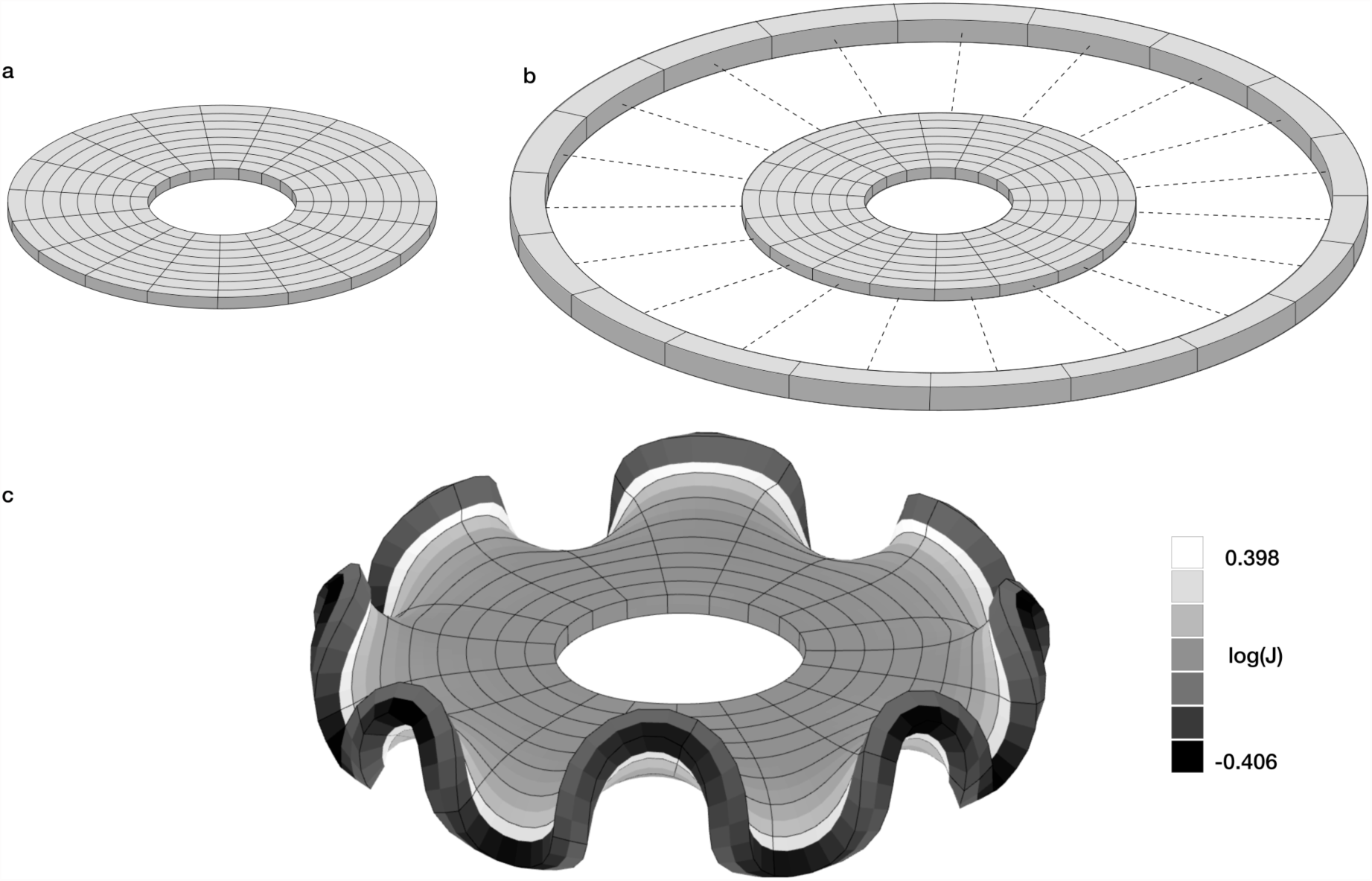
**Buckling induced by growth.** In this example, the outer most band of the ring is made to grow to twice its original size. This leads to the formation of wrinkles that minimise the elastic energy of the system. At equilibrium, there appears a heterogeneous pattern of stress, with residual stress gradients from the centre to the periphery, in angular bands and through the thickness of the ring. Homogeneous growth leads then to a heterogeneous pattern of residual stress at the microscopic scale, (a) Original configuration of the ring at rest, (b) After growth, the ring could reach a zero stress configuration only if its outer band were cut from the inner core. The dashed lines show the regions that should have to be stuck together to reconstruct the original ring, (c) The uncut ring after growth. The outer band is overall contracted, with gradients of residual stress from the inside to the outside of each fold. The immediately adjacent parts of the inner core are overall dilated, with a deformation that decreases as we approach the centre of the ring. Grey level: logarithm of the deformation (Jacobian of the deformation tensor). Black: log(J)=-0.406, white: log(J)=-0.398, bulk modulus *K* =100, shear modulus **μ**=100. An interactive simulation illustrating buckling induced by growth can be accessed at http://neuroanatomv.github.io/growth.

The first model of brain folding based on the emergence of a buckling instability was that of Richman et al (1975). This was a model of wrinkling where smooth, sinusoidal, folds developed because of a difference in growth and stiffness between the superior and inferior layers of the neocortex. The model proposed a process for producing folding through exclusively neocortical means (i.e., without requiring a cranial constraint). The model left unanswered, however, the questions of the constancy of species-specific folding patterns and the relationship between folding and neocortical organisation (cytoarchitecture, connectivity, function).

These questions were addressed by the model of Toro and Burnod (2005). In this 2D model, wrinkling was produced by the growth of a neocortical layer over an elastic and plastic substrate representing white matter and radial glia. In addition to elasticity, the tissues were also considered to be plastic: the heterogeneous mechanical stress produced by folding could induce local cortical growth and reabsorption, or the elongation and contraction of the white matter. Toro and Burnod (2005) observed that the formation and orientation of folds (wrinkles) could be modulated by cytoarchitectonic differences if they were already present, but also by the initial (unfolded) geometry of the cortical layer. This last mechanism could provide a purely geometric basis for the development and evolution of species-specific folding patterns (Toro, 2012).

The models by Richman et al (1975) and Toro and Burnod (2005) were exclusively theoretical, but more recent models based on mechanical parameters similar to those of real brains show that the morphology and stress patterns produced by buckling agree with those observed experimentally (Bayly et al., 2013; Budday et al., 2014a, 2014b; Tallinen et al., 2014).

These models suggest that mechanical instabilities induced by homogenous growth are sufficient to produce folds, without requiring specific, local, gyrogenetic processes such as a genetically determined, regionally higher rates of growth in pro-gyral regions (de Juan Romero et al., 2015; Lefèvre and Mangin, 2010; Reillo et al., 2011; Welker et al., 1990), specific cortico-cortical connectivity between pro-gyral walls (Van Essen, 1997; Hilgetag and Barbas, 2005, 2006), or a specific attachment of pro-sulcal regions (Régis et al., 2005; Smart and McSherry, 1986a, 1986b).

Additionally, these models suggest that the pattern of folding could be determined by the initial geometry of the neocortex. The orientation of folds in many species has been observed to correspond with the lines of principal curvature of the surface (Todd, 1982), which in many cases correspond with directions either concentric or orthogonal to the system formed by the corpus callosum and the Sylvian fissure (Régis et al., 2005; Toro and Burnod, 2003). In the same way that the geometry of a dome rigidifies its structure, the initial geometry of the neocortex makes some regions easier to fold than anothers, facilitating the development of folds in specific positions, with preferential orientations.

Previous theories explained the relationship between neocortical organisation and brain folding by encoding both of them in the genome. But if folding and folding patterns result from mechanical growth-induced buckling instabilities, we have to consider the possibility that, at least in part, **brain folding may have a causal effect on neocortical organisation**.

## 4. Effect of mechanical forces on neocortical development

The macroscopic, heterogeneous, residual stress induced by brain folding could regionally modulate cell proliferation, cell fate and cell shape, influence axonal guidance and even synaptic activity, thus affecting the organisation of the neocortex (Franze, 2013). Localised stress is known to modulate tissue volume, with tension facilitating tissue growth and compression promoting reabsorption (Fung, 1993; Rodriguez et al., 1994). Compressing a tumour spheroid formed from carcinoma cells decreases cell proliferation in the regions of high mechanical stress, and cells could even undergo apoptosis if stress is sufficiently high (Cheng et al., 2009; Montel et al., 2012). In neural stem cells culture, the rate of cell division reaches a peak at 1-4 kPa, but decreases in stiffer or softer substrates. In gels softer than ∼10 Pa, the spreading, self-renewal and differentiation of neural stem cells is almost completely inhibited (Saha et al., 2008). Mechanical stress can affect progenitor cells directly, but also indirectly, by affecting the structure of the extra-cellular matrix. Mechanical stress could result in differential local concentrations of extra-cellular matrix components, which in turn could lead to differences in cell signalling and adhesion (Lutolf et al., 2009).

In addition to cell proliferation, cell fate and differentiation can be regulated by mechanical forces. Engler et al (2006) showed that naive mesenchymal stem cells cultured on a substrate mimicking the elasticity of brain, muscle or bone, differentiated respectively into branched cells similar to primary neurones, spindle-shaped cells similar to myoblasts or polygonal cells similar to osteoblasts. Interestingly, whereas reprogramming these lineages by the addition of soluble induction factors was possible only during the initial week in culture, changes in cell fate through manipulation of matrix elasticity were possible even after several weeks. In neural stem cells culture, Saha et al (2008) showed that soft gels (∼100-500 Pa) produced mostly neurones, whereas increasingly harder gels (up to 1-10 kPa) produced progressively more glial cells. In vivo, neocortical neurones grow along glial cells, which may be due in part to the fact that glia are significantly softer than their neighbouring neurones (Lu et al., 2006).

The shape of differentiated cells can also respond to the mechanical properties of their environment. Neurones cultured on soft substrates (50-300 Pa) develop up to 3 times more branching than those cultured on stiffer gels (300-550 Pa, Flanagan et al (2002)). Neurones also seem to require a certain amount of mechanical tension to mature. In vitro, the axonal branches which attach strongly to the substrate are conserved and stabilised, whereas the remaining branches are retracted or eliminated (Anava et al., 2009). Pulling axons at different speeds not only increases their length accordingly, but also their calibre (Bray, 1984). That is, nerve cell respond to tension applied along their axons by building more axon, increasing the length and number of its microtubules, neurofilaments and membrane components.

Buckling models of neocortical folding predict gradients of mechanical stress between gyri and sulci spanning the thickness of the cortex and close white matter (such as those in Fig. 3c). These gradients could participate to the establishment of the differences in cell shape and layering observed in vertebrates (Welker et al., 1990). Cell migration and axonal pathfinding respond to mechanical clues (Saez et al., 2007), and could be guided by the mechanical stiffness gradients produced by growth-driven buckling instabilities. Finally, tension in neurones has been observed to produce an active accumulation of the synaptic vesicles involved in neurotransmitter releasing to postsynaptic cells (Ahmed et al., 2012), which leads to a possible modulation of the synaptic activity by the mechanical properties of the surrounding tissue.

## 5. Discussion

### 5.1. A hypothesis about neocortical development and evolution

From the previous considerations emerges a model where the development and evolution of the neocortex result from the interaction of genetic, mechanical and activity-driven processes. Brain development is the plastic deployment of a complex organisation, involving a massive production of cells, their migration, differentiation, specialisation and interconnection. Under the forces of biological growth, brain tissue undergoes major changes in geometry correlated with constant changes in the distribution of mechanical stress. This is the context within which patterning centres diffuse morphogenes and signalling molecules. Through their effect on transcription factors and cell behaviour (proliferation, differentiation,…), these molecules can affect local growth and the mechanical properties of the developing tissue, changing the global distribution of stress. In return, mechanical gradients can affect gene expression and cell behaviour back, in a continuous interplay between biological and physical forces. Neocortical organisation is layed out in this dynamic substrate, under the influence of gradients of morphogenes, signalling molecules and mechanical stress.

New cortical areas could develop because of an increase in the complexity of the intrinsic response to transcription factors. Morphogenetic gradients established by the diffusion of molecular signals from patterning centres could provide unique coordinates for each point in the cortical mantle. This could regulate the local expression of different transcription factors. In response to different combinations of transcription factor levels, cortical tissue could develop a restricted number of cell identities: the basis of the future cortical areas. Through these morphogenetic gradients, intrinsic genetic processes could also regulate the expression of axonal guidance molecules, and control the formation of specific connectivity patterns.

But new areas could also develop because of mechanical morphogenesis. Homogeneous cortical growth could induce buckling instabilities that could produce cortical folding. Each fold would be characterised by a geometric deformation – a gyrus – and a mechanical stress pattern, with tension in gyral crowns, compression in sulcal fundi, and a gradual change in tension across cortical layers and close white matter. For comparable thickness, a large cerebral cortex should develop more folds, i.e., more of these discontinuous mechanical domains. The geometric and mechanical changes within each fold would introduce localised changes in cell identity and influence the formation of brain connectivity. Depending on their spatial situation and time of development, neurones within each fold would establish different sets of connections with the rest of the brain, in particular with sensory and motor thalamic nuclei.

The causes of the formation of a new area could be then genetic – a change in the complexity of the response to gradients of transcription factors – or mechanical – a change in the number of mechanical domains. In both cases, whether new cortical areas appear because of a change in the genetic program or because of the effect if mechanical morphogenesis, the organisation produced by intrinsic genetic or mechanical factors could be plastically modified by neuronal activity, which in turn could influence gene expression or change the mechanical properties of the tissue.

### 5.1. Consequences of mechanical morphogenesis

Across species, the number of cortical areas and the complexity of the connnectivity patterns increases with brain size (Bourgeois, 1997; Krubitzer, 2007). In a small brain, the number of cortical areas should be determined by the discrete number of possible responses of the nervous tissue to different levels of transcription factors, by the response to monotonous gradients of mechanical stress (that should be present even in the absence of buckling and brain folding), or by the plastic response to activity driven by thalamic afferents. An evolutionary change in brain size sufficient to induce cortical folding should lead to the creation of new cortical areas and connectivity patterns even if the number of responses to transcription factor levels or to neuronal activity stays the same. Because large brains are progressively more folded than small ones, and develop an exponentially larger amount of connections, mechanical morphogenesis could explain in part the augmentation in the number of cortical areas.

The effect of mechanical morphogenesis on neocortical organisation could also be studied within a single species, where genetic differences are much smaller, and restricted mostly to genetic polymorphism. Despite a high genetic similarity (Kaessmann et al., 2001), humans exhibit a remarkable diversity in total brain volume, mostly due to differences in total cortical surface area. Large human brains have disproportionately more cortical surface area and are significantly more folded than small ones (Germanaud et al., 2012; Toro et al., 2008). These differences are mainly due to tertiary folds (shallow, late developing and variable) and in some cases to secondary folds, but the deep, early developing folds, are very stable. Nevertheless, we should observe a larger number of neocortical areas in larger, more folded brains, as well as differences in the connectivity patterns. Some reports suggest indeed that the presence of supplementary folds among humans, such as the paracingulate fold of the cingulate cortex, can be associated with differences in behaviour (Fornito et al., 2004, 2006) and cytoarchitecture (Vogt et al., 1995). In addition to natural inter-subject diversity, pathologies such as lissencephaly and microgyria, should provide an opportunity to study the effect of mechanical morphogenesis on neocortical organisation. We should observe a change in the number of cortical areas and connectivity patterns associated with the changes in cortical folding, especially if the pattern of early developing primary folds is affected.

### 5.2. Experimental perturbations of mechanical morphogenesis

If our hypothesis is correct, mechanical perturbations of the developing cortex should be able to modify the area map, add new areas, or produce new connectivities. As with genetic perturbations (gene knockouts, knockins, etc.) or perturbations of neuronal activity (sensory deprivation, rewiring, etc.), perturbations of mechanical morphogenesis will require to develop appropriate controls to ensure that the effects observed are due to the intended perturbation and not to experimental artefacts.

The ferret has been often used to study brain folding. At birth, its neocortex is completely smooth, but after a first stage of growth without folding, somewhere between P4 and P8 it starts to fold. After one month, the ferret has a richly folded neocortex with deep sulci organised in a characteristic folding pattern. If brain folding is produced because of a buckling instability, the ferret neocortex should be at a maximally unstable state right before folding, probably by P4. A small mechanical perturbation introduced at this time should be enough to trigger the formation of an artificial fold. Mechanically, the most probable orientation of a fold should be along one of the 2 principal curvature directions of the surface before folding (Tallinen et al., 2014; Todd, 1982; Toro and Burnod, 2003). If a natural fold is formed along one principal curvature direction, it should be possible in theory to force the formation of an artificial fold along the other, orthogonal, principal curvature direction.

In wild-type ferrets, an increased rate of neurogenesis (measured by a higher density of *Tbr2+* progenitors) has been observed in regions that will later underlie a gyrus (de Juan Romero et al., 2015; Reillo et al., 2011). If neurogenesis is influenced by the mechanical stress that builds a fold, we should observe a new, ectopic increase of neurogenesis under the artificial gyri, or a decrease under the artificial sulci. As a result, the orientation of the cortical areas in the mechanically perturbed cortex should be different than in the wild-type ferret, and correspond with the orientation of the new folding pattern. Additionally, if connectivity is influenced by the geometry of folding, we should observe an altered pattern of cortico-cortical and cortico-subcortical connectivity. Neuronal activity should further influence the organisation of the new cortical areas, depending on the sensory modalities of the thalamic afferents they receive. The behaviour of the animals should also be changed as a result of the new brain organisation. In ferrets, auditory fear conditioning is faster than visual fear conditioning, likely because of the more direct connection from the auditory thalamic nuclei to the amygdala than from the visual thalamic nuclei to the amygdala (Newton et al., 2004). In rewired ferrets, however, visual fear conditioning is mediated by auditory nuclei and is then faster than in control animals. The same phenomenon should be observed in ferrets with artificial folds: the different patterns of connections between the new cortical areas should produce similar behavioural alterations.

In conclusion, we have outlined a hypothesis on the role of mechanical morphogenetic processes in the definition of neocortical organisation. If this hypothesis is correct, brain folding should have a causal effect on brain development and play, together with genetic and activity-dependent processes, an important role on the evolution of the neocortex.

## Acknowledgments

We would like to thank J-P Bourgeois, T. Bourgeron, E. Cerda, L. Mahadevan, L. Ronan and M. Trejo for helpful discussions.

## References

Ahmed, W.W., Li, T.C., Rubakhin, S.S., Chiba, A., Sweedler, J.V., and Saif, T.A. (2012). Mechanical Tension Modulates Local and Global Vesicle Dynamics in Neurons. Cell. Mol. Bioeng. 5, 155–164.

Amar, M.B., Müller, M.M., and Trejo, M. (2012). Petal shapes of sympetalous flowers: the interplay between growth, geometry and elasticity. New J. Phys. 14, 085014.

Anava, S., Greenbaum, A., Jacob, E.B., Hanein, Y., and Ayali, A. (2009). The Regulative Role of Neurite Mechanical Tension in Network Development. Biophys. J. 96, 1661–1670.

Bayly, P.V., Okamoto, R.J., Xu, G., Shi, Y., and Taber, L.A. (2). A cortical folding model incorporating stress-dependent growth explains gyral wavelengths and stress patterns in the developing brain. Phys. Biol. 10, 016005.

Bishop, K.M., Goudreau, G., and O’Leary, D.D. (2000). Regulation of area identity in the mammalian neocortex by Emx2 and Pax6. Science 288, 344–349.

Borrell, V., and Götz, M. (2014). Role of radial glial cells in cerebral cortex folding. Curr. Opin. Neurobiol. 27, 39–46.

Bourgeois, J. (1997). Synaptogenesis, heterochrony and epigenesis in the mammalian neocortex. Acta Paediatr. 86, 27–33.

Bowden, N., Brittain, S., Evans, A.G., Hutchinson, J.W., and Whitesides, G.M. (1998). Spontaneous formation of ordered structures in thin films of metals supported on an elastomeric polymer. Nature 393, 146–149.

Bray, D. (1984). Axonal growth in response to experimentally applied mechanical tension. Dev. Biol. 102, 379–389.

Brodmann, K. (1909). Vergleichende Lokalisationslehre der Grosshimrinde (Leipzig: Verlag von Johann Ambrosias Barth).

Brodmann, K., and Gary, L.J. (2006). Brodmann’s localization in the cerebral cortex the principles of comparative localisation in the cerebral cortex based on cytoarchitectonics ([New York, NY]: Springer).

Budday, S., Raybaud, C., and Kuhl, E. (7). A mechanical model predicts morphological abnormalities in the developing human brain. Sci. Rep. 4.

Budday, S., Steinmann, P., and Kuhl, E. (2014). The role of mechanics during brain development. J. Mech. Phys. Solids 72, 75–92.

Cerda, E., and Mahadevan, L. (2003). Geometry and Physics of Wrinkling. 1–5.

Cheng, G., Tse, J., Jain, R.K., and Munn, L.L. (2009). Micro-environmental mechanical stress controls tumor spheroid size and morphology by suppressing proliferation and inducing apoptosis in cancer cells. PloS One 4, e4632.

Ciarletta, P. (2013). Buckling Instability in Growing Tumor Spheroids. Phys. Rev. Lett. 110, 158102.

Constantine-Paton, M., and Law, M.I. (1978). Eye-specific termination bands in tecta of three-eyed frogs. Science 202, 639–641.

Davidovitch, B., Schroll, R.D., Vella, D., Adda-Bedia, M., and Cerda, E.A. (2011). Prototypical model for tensional wrinkling in thin sheets. Proc. Natl. Acad. Sci. U. S. A. 108, 18227–18232.

Engler, A.J., Sen, S., Sweeney, H.L., and Discher, D.E. (2006). Matrix elasticity directs stem cell lineage specification. Cell 126, 677–689.

Espinosa, J.S., and Stryker, M.P. (2012). Development and Plasticity of the Primary Visual Cortex. Neuron 75, 230–249.

Van Essen, D.C. (1997). A tension-based theory of morphogenesis and compact wiring in the central nervous system. Nature 385, 313–318.

Fischl, B. (2013). Estimating the Location of Brodmann Areas from Cortical Folding Patterns Using Histology and Ex Vivo MRI. In Microstructural Parcellation of the Human Cerebral Cortex, S. Geyer, and R. Turner, eds. (Springer Berlin Heidelberg), pp. 129–156.

Fischl, B., Rajendran, N., Busa, E., Augustinack, J., Hinds, O., Yeo, B.T.T., Mohlberg, H., Amunts, K., and Zilles, K. (2008). Cortical Folding Patterns and Predicting Cytoarchitecture. Cereb. Cortex 18, 1973–1980.

Flanagan, L.A., Ju, Y.-E., Marg, B., Osterfield, M., and Janmey, P.A. (2002). Neurite branching on deformable substrates. Neuroreport 13, 2411–2415.

Fornito, A., Yücel, M., Wood, S., Stuart, G.W., Buchanan, J.-A., Proffitt, T., Anderson, V., Velakoulis, D., and Pantelis, C. (2004). Individual differences in anterior cingulate/paracingulate morphology are related to executive functions in healthy males. Cereb. Cortex N. Y. N 1991 14, 424–431.

Fornito, A., Yücel, M., Wood, S.J., Proffitt, T., McGorry, P.D., Velakoulis, D., and Pantelis, C. (2006). Morphology of the paracingulate sulcus and executive cognition in schizophrenia. Schizophr. Res. 88, 192–197.

Franze, K. (2013). The mechanical control of nervous system development. Dev. Camb. Engl. 140, 3069–3077.

Fukuchi-Shimogori, T., and Grove, E. a (2001). Neocortex patterning by the secreted signaling molecule FGF8. Science 294, 1071–1074.

Fung, Y.C. (1993). Biomechanics: mechanical properties of living tissues (New York: Springer-Verlag).

Geng, G., Johnston, L. a, Yan, E., Britto, J.M., Smith, D.W., Walker, D.W., and Egan, G.F. (2009). Biomechanisms for modelling cerebral cortical folding. Med. Image Anal. 13, 920–930.

Germanaud, D., Lefèvre, J., Toro, R., Fischer, C., Dubois, J., Hertz-Pannier, L., and Mangin, J.-F. (2012). Larger is twistier: Spectral analysis of gyrification (SPANGY) applied to adult brain size polymorphism. NeuroImage 63, 1257–1272.

Hamasaki, T., Leingärtner, A., Ringstedt, T., and O’Leary, D.D.M. (2004). EMX2 regulates sizes and positioning of the primary sensory and motor areas in neocortex by direct specification of cortical progenitors. Neuron 43, 359–372.

Hannezo, E., Prost, J., and Joanny, J.-F. (2011). Instabilities of monolayered epithelia: shape and structure of villi and crypts. Phys. Rev. Lett. 107, 078104.

Hansen, D.V., Lui, J.H., Parker, P.R.L., and Kriegstein, A.R. (2010). Neurogenic radial glia in the outer subventricular zone of human neocortex. Nature 464, 554–561.

Hilgetag, C.C., and Barbas, H. (2005). Developmental mechanics of the primate cerebral cortex. Anat. Embryol. (Berl.) 210, 411–417.

Hilgetag, C.C., and Barbas, H. (2006). Role of mechanical factors in the morphology of the primate cerebral cortex. PloS Comput. Biol. 2, e22–e22.

Hohlfeld, E., and Mahadevan, L. (2011). Unfolding the Sulcus. Phys. Rev. Lett. 106.

De Juan Romero, C., Bruder, C., Tomasello, U., Sanz-Anquela, J.M., and Borrell, V. (2015). Discrete domains of gene expression in germinal layers distinguish the development of gyrencephaly. EMBO J.

Kaas, J.H. (2012). Evolution of columns, modules, and domains in the neocortex of primates. Proc. Natl. Acad. Sci. 109, 10655–10660.

Kaessmann, H., Wiebe, V., Weiss, G., and Pääbo, S. (2001). Great ape DNA sequences reveal a reduced diversity and an expansion in humans. Nat. Genet. 27, 155–156.

Krubitzer, L. (2007). The magnificent compromise: cortical field evolution in mammals. Neuron 56, 201–208.

Krubitzer, L. (2009). In search of a unifying theory of complex brain evolution. Ann. N. Y. Acad. Sci. 1156, 44–67.

Lefèvre, J., and Mangin, J.-F. (2010). A reaction-diffusion model of human brain development. PloS Comput. Biol. 6, e1000749–e1000749.

Lu, Y.-B., Franze, K., Seifert, G., Steinhauser, C., Kirchhoff, F., Wolburg, H., Guck, J., Janmey, P., Wei, E.- Q., Kas, J., et al. (2006). Viscoelastic properties of individual glial cells and neurons in the CNS. Proc. Natl. Acad. Sci. 103, 17759–17764.

Lutolf, M.P., Gilbert, P.M., and Blau, H.M. (2009). Designing materials to direct stem-cell fate. Nature 462, 433–441.

Mallamaci, A., Muzio, L., Chan, C.H., Parnavelas, J., and Boncinelli, E. (2000). Area identity shifts in the early cerebral cortex of Emx2-/- mutant mice. Nat. Neurosci. 3, 679–686.

Von Melchner, L., Pallas, S.L., and Sur, M. (2000). Visual behaviour mediated by retinal projections directed to the auditory pathway. Nature 404, 871–876.

Merlin, S., Horng, S., Marotte, L.R., Sur, M., Sawatari, A., and Leamey, C.A. (2013). Deletion of Ten-m3 Induces the Formation of Eye Dominance Domains in Mouse Visual Cortex. Cereb. Cortex 23, 763–774.

Montel, F., Delarue, M., Elgeti, J., Vignjevic, D., Cappello, G., and Prost, J. (2012). Isotropic stress reduces cell proliferation in tumor spheroids. New J. Phys. 14, 055008.

Newton, J.R., Ellsworth, C., Miyakawa, T., Tonegawa, S., and Sur, M. (2004). Acceleration of visually cued conditioned fear through the auditory pathway. Nat. Neurosci. 7, 968–973.

Nonaka-Kinoshita, M., Reillo, I., Artegiani, B., Àngeles Martínez-Martínez, M., Nelson, M., Borrell, V., and Calegari, F. (2013). Regulation of cerebral cortex size and folding by expansion of basal progenitors. EMBO J. 32, 1817–1828.

O’Leary, D.D.M. (1989). Do cortical areas emerge from a protocortex? Trends Neurosci. 12, 400–406.

O’Leary, D.D.M., Stocker, A.M., Zembrzycki, A., and Rakic, J.L.R.R. (2013). Chapter 4 - Area Patterning of the Mammalian Cortex. In Patterning and Cell Type Specification in the Developing CNS and PNS, (Oxford: Academic Press), pp. 61–85.

Pocivavsek, L., Dellsy, R., Kern, A., Johnson, S., Lin, B., Lee, K.Y.C., and Cerda, E. (2008). Stress and Fold Localization in Thin Elastic Membranes. Science 320, 912–916.

Puelles, L. (2009). Forebrain Development: Prosomere Model. In Encyclopedia of Neuroscience, (Elsevier), pp. 315–319.

Puelles, L. (2013). Chapter 10 - Plan of the Developing Vertebrate Nervous System: Relating Embryology to the Adult Nervous System (Prosomere Model, Overview of Brain Organization). In Patterning and Cell Type Specification in the Developing Cns and Pns, J.L.R.R. Rakic, ed. (Oxford: Academic Press), pp. 187–209.

Rakic, P. (1988). Specification of cerebral cortical areas. Science 241, 170–176.

Rakic, P., Ayoub, A.E., Breunig, J.J., and Dominguez, M.H. (2009). Decision by division: making cortical maps. Trends Neurosci. 32, 291–301.

Régis, J., Mangin, J.-F., Ochiai, T., Frouin, V., Riviére, D., Cachia, A., Tamura, M., and Samson, Y. (2005). “Sulcal root” generic model: a hypothesis to overcome the variability of the human cortex folding patterns. Neurol. Med. Chir. (Tokyo) 45, 1–17.

Reillo, I., de Juan Romero, C., Garcia-Cabezas, M.A., and Borrell, V. (2011). A Role for Intermediate Radial Glia in the Tangential Expansion of the Mammalian Cerebral Cortex. Cereb. Cortex 21, 1674–1694.

Richman, D.P., Stewart, R.M., Hutchinson, J.W., and Caviness, V.S. (1975). Mechanical model of brain convolutional development. Science 189, 18–21.

Rivlin, R.S. (1948). Large Elastic Deformations of Isotropic Materials. IV. Further Developments of the General Theory. Philos. Trans. R. Soc. Math. Phys. Eng. Sci. 241, 379–397.

Rodriguez, E.K., Hoger, A., and McCulloch, A.D. (1994). Stress-dependent finite growth in soft elastic tissues. J. Biomech. 27, 455–467.

Roe, A.W., Pallas, S.L., Hahm, J.O., and Sur, M. (1990). A map of visual space induced in primary auditory cortex. Science 250, 818–820.

Roe, A.W., Pallas, S.L., Kwon, Y.H., and Sur, M. (1992). Visual projections routed to the auditory pathway in ferrets: receptive fields of visual neurons in primary auditory cortex. J. Neurosci. Off. J. Soc. Neurosci. 12, 3651–3664.

Ronan, L., and Fletcher, P.C. (2014). From genes to folds: a review of cortical gyrification theory. Brain Struct. Amp Funct.

Saez, A., Ghibaudo, M., Buguin, A., Silberzan, P., and Ladoux, B. (2007). Rigidity-driven growth and migration of epithelial cells on microstructured anisotropic substrates. Proc. Natl. Acad. Sci. 104, 8281–8286.

Saha, K., Keung, A.J., Irwin, E.F., Li, Y., Little, L., Schaffer, D.V., and Healy, K.E. (2008). Substrate Modulus Directs Neural Stem Cell Behavior. Biophys. J. 95, 4426–4438.

Sharma, J., Angelucci, A., and Sur, M. (2000). Induction of visual orientation modules in auditory cortex. Nature 404, 841–847.

Smart, I.H., and McSherry, G.M. (1986a). Gyrus formation in the cerebral cortex in the ferret. I. Description of the external changes. J. Anat. 146, 141–152.

Smart, I.H., and McSherry, G.M. (1986b). Gyrus formation in the cerebral cortex of the ferret. II. Description of the internal histological changes. J. Anat. 147, 27–43.

Stahl, R., Walcher, T., De Juan Romero, C., Pilz, G.A., Cappello, S., Irmler, M., Sanz-Aquela, J.M., Beckers, J., Blum, R., Borrell, V., et al. (2013). Trnp1 Regulates Expansion and Folding of the Mammalian Cerebral Cortex by Control of Radial Glial Fate. Cell 153, 535–549.

Sur, M., Garraghty, P., and Roe, A. (1988). Experimentally induced visual projections into auditory thalamus and cortex. Science 242, 1437–1441.

Tallinen, T., Chung, J.Y., Biggins, J.S., and Mahadevan, L. (2014). Gyrification from constrained cortical expansion. Proc. Natl. Acad. Sci. U. S. A. 111, 12667–12672.

Todd, P.H. (1982). A geometric model for the cortical folding pattern of simple folded brains. J. Theor. Biol. 97, 529–538.

Toro, R. (2012). On the Possible Shapes of the Brain.

Toro, R., and Burnod, Y. (2003). Geometric atlas: modeling the cortex as an organized surface. NeuroImage 20, 1468–1484.

Toro, R., and Burnod, Y. (2005). A morphogenetic model for the development of cortical convolutions. Cereb Cortex 15, 1900–1913.

Toro, R., Perron, M., Pike, B., Richer, L., Veillette, S., Pausova, Z., and Paus, T. (2008). Brain size and folding of the human cerebral cortex. Cereb. Cortex N. Y. N 1991 18, 2352–2357.

Vogt, B.A., Nimchinsky, E.A., Vogt, L.J., and Hof, P.R. (1995). Human cingulate cortex: surface features, flat maps, and cytoarchitecture. J. Comp. Neurol. 359, 490–506.

Welker, W.I., and Campos, G.B. (1963). Physiological significance of sulci in somatic sensory cerebral cortex in mammals of the family procyonidae. J. Comp. Neurol. 120, 19–36.

Welker, W., Jones, E.G., and Peters, A. (1990). Why Does Cerebral Cortex Fissure and Fold? (Boston, MA: Springer US), pp. 3–136.

Xie, W.-H., Li, B., Cao, Y.-P., and Feng, X.-Q. (2014). Effects of internal pressure and surface tension on the growth-induced wrinkling of mucosae. J. Mech. Behav. Biomed. Mater. 29, 594–601.

Xu, G., Bayly, P.V., and Taber, L. a (2009). Residual stress in the adult mouse brain. Biomech. Model. Mechanobiol. 8, 253–262.

Xu, G., Knutsen, A.K., Dikranian, K., Kroenke, C.D., Bayly, P.V., and Taber, L. a (2010). Axons pull on the brain, but tension does not drive cortical folding. J. Biomech. Eng. 132, 071013–071013.

